# Climate change triggers morphological and life-history evolution in response to predators

**DOI:** 10.1101/001263

**Authors:** Edmund M. Hart, Nicholas J. Gotelli

## Abstract

Although climate change is expected to reorganize entire communities, this restructuring might reflect either direct ecological or evolutionary responses to abiotic conditions or indirect effects mediated through altered species interactions. We tested the hypothesis that changes in trophic interaction strength due to altered predator abundance have a cascading evolutionary response in a prey species (*Daphnia pulex*). Using a multiyear / multigenerational field experiment, we manipulated 12 open aquatic mesocosms to simulate hydrological conditions under climate change. After a three-year press manipulation, we collected *Daphnia pulex* from each pond and raised them in a common garden. Using quantitative genetic methods, we measured a series of quantitative traits every other day on 108 individuals for eight weeks. There was a significant decrease in tail spine length and population growth rate in groups exposed to the most extreme future climate scenarios. Structural equation models demonstrated that trait changes were best explained as an indirect effect of climate change treatments mediated through changes in predator abundance. Our results suggest climate change can trigger a cascade of ecological and evolutionary forces by reducing predator density, which in turn acts as a selective force leading to evolutionary change in prey morphology and life history.

## Introduction

Global climate change is expected to lead to major shifts in the composition of communities and the structure of populations (Tylianakis et al. 2008, Kratina et al. 2012). These shifts often occur because of individualistic species responses to direct abiotic factors such as temperature (Menge et al. 2008) or precipitation (Hart and Gotelli 2011). The resulting assemblage may exhibit altered trophic structure (Urban et al. 2012b) which could trigger additional change. What remains unclear is how the direct responses of one species to climate change lead to indirect effects on other community members, and whether these effects are strictly ecological or have an evolutionary basis. Complex community-level shifts in response to climate change are themselves potential selective agents (Harmon et al. 2009) and could trigger evolutionary responses in a focal species.

Direct responses to abiotic factors associated with climate change are important (Lavergne et al. 2010, Hoffmann and Sgro 2011), and there are well-documented examples of direct evolutionary responses of populations to elevated CO_2_ (Collins and Bell 2004), temperature (Jump et al. 2008), and pH (Lohbeck et al. 2012). However, populations may have complex responses to climate change that are driven by direct changes in abiotic factors (e.g. temperature, precipitation) and by indirect changes in the density of predators or competitors (Harmon et al. 2009, Urban et al. 2012a). For example laboratory populations of *Daphnia magna* evolved differently in response to increased temperature in isolated populations versus those that were exposed to an entire assemblage of competitors and predators (Van Doorslaer et al. 2010).

Although terrestrial communities have been the primary research focus, climate change will also have important impacts in aquatic systems (Kratina et al. 2012). Climate change is expected to lead not only to increases in air temperature, but also to changes in precipitation intensity and drought stress (Frumhoff et al. 2007). Freshwater food webs will be strongly affected by the resulting changes in hydrology, particularly in ephemeral habitats such as vernal ponds (Brooks 2009). One key taxon in many aquatic food webs is *Daphnia pulex* (Branchiopoda: Cladocera) a filter-feeding crustacean that is abundant in many temporary and permanent aquatic habitats (Lynch 1980). In laboratory studies, *Daphnia* life history and morphology respond directly to changes in temperature (VanDoorslaer et al. 2010), as well as to the presence of predators, which can cause rapid evolution in growth and spine length (Spitze 1991, Fisk et al. 2007).

Here we present the results of a three year field experiment in which simulated changes in the hydrology of vernal ponds reduce the abundance of aquatic predators, leading to evolutionary change over several generations in the morphology and life history of *Daphnia pulex*. Using a multi-generation field study and a common garden experiment, we show that increased evapotranspiration and drought stress, both predicted with climate change (Brooks 2009), reduce the abundance of invertebrate predators, which triggers rapid evolutionary shifts in the spine length and intrinsic rate of increase of *D. pulex.* Our results demonstrate that climate change can initiate a cascading response by reducing the abundance of predators therefore altering trophic interaction strength, which leads to evolutionary change in the morphology and life history of prey populations.

## Materials and Methods

### Study organism and system

*Daphnia pulex* (Branchiopoda: Cladocera) is a filter-feeding zooplankter that consumes bacteria and phytoplankton and is common in a wide variety of temporary and permanent aquatic habitats. *D. pulex* individuals usually survive for 16 instars (Lynch 1980) and exhibit a sigmoid growth curve for body size, with rapid growth to first reproduction between the fourth and sixth instars, and then slower growth in later instars as body size approaches an asymptote (Green 1956, Lynch 1980). In our system, *D. pulex* is obligately parthenogenetic (Hebert and Finston 2001), with mature diploid females clonally producing daughters every instar (Green 1956).

Female body size, clutch size, and neonate size are all correlated within isofemale lines, but there is substantial variation in traits among clones (Green 1956). Because *D. pulex* undergoes frequent cycles of clonal reproduction, differential survival of clonal lineages can lead to highly adapted local populations (Haag et al. 2006).

### Effects of climate change on vernal ponds

Vernal ponds are fishless habitats that fill in the spring (vernal) or fall (autumnal) and hold water for at least 4 months, but are seasonally dry. Regional climate change models predict a warming of 3.5^o^ to 6.5^o^ F by 2100 under low-emission scenarios and an even larger increase under high-emission scenarios (Frumhoff et al. 2007). Climate change scenarios for the northeastern United States also predict an increased water budget in the winter and spring and increased deficit in the summer and fall (Frumhoff et al. 2007). Precipitation events are likely to become more variable, with longer periods of drought followed by more intense deluges (Hayhoe et al. 2008). Two effects of climate change will alter the hydroperiod of vernal ponds: increased evapotranspiration, and more variable precipitation. More variable spring and summer rainfall combined with greater evapotranspiration will increase the drying frequency and the variance of water level (Brooks 2009).

### Experimental design

In May of 2008, we used a small excavator to dig 81 ponds within a 500 × 500 m area at the University of Vermont Jericho Research Forest (44.45° N, 73.00° W). Ponds were dispersed and constructed along a network of logging roads to facilitate access and reduce spatial autocorrelation. We cleared ponds of roots and rocks and smoothed them into shape using hand tools until they were approximately 1.5 m in diameter and 60 cm deep. We lined each pond with 3 × 3 m sheets of 0.254 mm thick black plastic, covered the bottom of the ponds with a layer of dirt and leaf litter, and filled them with water from an on-site spring-fed well. The final depth of the full ponds was approximately 45 cm, with a starting volume of approximately 5000 L. We inoculated each pond on 16 June 2008 with 1L of water from a mixture of 81 plankton tows and 243 dip-net samples collected from a large local vernal pond. The average number of *Daphnia pulex* that was introduced into each pond was ∼150 individuals.

We simultaneously varied the rate at which a pond lost water (water loss rate), and the frequency and intensity of rainfall events (drought severity) to experimentally generate a range of hydrology profiles that represented current to future climates. Our experiment had nine levels each of water loss rate and drought severity applied to 81 artificial ponds in a fully-crossed response-surface design. Our treatments mimicked two aspects of projected climate change: increased evapotranspiration (Yu et al. 2002) and increased variability in precipitation (Hayhoe et al. 2008). We simulated a model of linear rate of water loss, allowing ponds to hold water from between 50 and 180 days, to mimic changes in evapotranspiration (Yu et al. 2002). Each treatment level represented an increase in pond hydroperiod of 15 days. This factor combined evaporation from increased temperature (Hayhoe et al. 2008) and increased water usage by trees (transpiration; Bonan 2008). The precipitation treatment was a single parameter with two levels tied together: as rainfall probability decreased, rainfall intensity increased. We estimated rainfall probabilities from the past 51 years of rainfall data (April to September from 1957 to 2007) recorded at Burlington International Airport (44.47° N, −73.15°). The mean daily probability of rainfall over those 6 months was 0.39, with a minimum of 0.18 and a maximum of 0.53. We created nine treatment levels that ranged evenly between 0.4 (the 51-year mean) and 0.03 (high potential for an extreme drought). Levels 1 – 5 were within current norms, and the remaining 4 levels represented extreme drought frequencies that Vermont has not yet experienced. Tied to each rainfall probability was a probability of an intense rainfall event that ranged from 0.01 to 0.1. An intense rainfall event was defined as any event in the 95th percentile of all rainfall events since 1957. Using the two parameters of water loss rate and drought severity, we created a simple model of vernal pond hydrology:

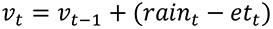

The pond volume *v* at time *t* is the volume at the previous time step plus rainfall (*rain_t_*, determined by drought severity) minus evapotranspiration (*et_t_*, determined by the water loss rate). We calculated the initial volume of each pond as one half of an ovoid sphere with a short radius of 42 cm and a long radius of 150 cm. The model was used to calculate the daily water balance by first randomly determining whether it would rain that day with the probability *p_i_* of rainfall in drought severity level *i.* If the algorithm specified rain, the rainfall event was characterized as an intense storm event with probability *s_i_.* Depending on the outcome, the model drew randomly from a gamma distribution fit to the past 51 years of data for whichever month the simulation was in (Apr – Sept). If an extreme rainfall event was selected in the model, then the amount was specified by a random draw from the distribution of previous extreme events. Thus, intense rainfall events were based on a statistical distribution, but we simulated future climates by making intense rainfall events more frequent.

### Common garden design

After three years of treatment application, we sampled the most extreme combinations of the experimental parameter space to create a 2 × 2 factorial ANOVA design. Although the ponds were not identical replicates, the hydrological profiles within the 4 clusters were very similar to one another. We collected 20 individual *Daphnia pulex* from each pond in late August 2010 from dip-net sweeps placed in a collection tray and live collected with an eye dropper. Field collected individuals were isolated in 250 ml glass jars. We established 240 isofemale lines, raised in filtered water that we changed every 5 days. All isofemales were raised in a Percival growth chamber on a 14:10 day:night cycle and a 23:18 day:night temperature regime (Spitze 1993).

In order to minimize maternal effects (Bernardo 1996), we raised lines over several generations in the growth chamber before beginning the life history and growth measurements on January 1^st^ 2011. Using a spectrophotometer, we diluted a stock of live *Nannochloropsis* (green algae, Carolina Biological Supply) food solution to 4 mg C/L and fed the isofemale lines every other day; this feeding regime ensured there was no reduction in fecundity due to food limitation (Lampert 1978). We randomly selected 3 isofemale lines per pond from the available 20 and isolated three offspring per isofemale line from their first clutch (3 clones / isofemale × 3 isofemales / pond × 3 ponds / treatment × 2 levels of drying rate × 2 levels of rainfall = 108 replicates) and photographed individuals every other day to measure morphological characters. We measured clutch size by counting live-born offspring in each jar, after which we removed them. We recorded all life history data on these individuals. Because all individuals were raised through multiple generations in a common garden, differences in average measured traits should reflect genetic differences among populations, not maternal (Bernardo 1996) or early-environment effects (Spitze 1991, Conner and Hartl 2004, Hansen et al. 2012). The common garden experiment ran from 1 January to 14 February 2011, and we measured a total of 251 individuals. If an individual did not survive to produce 3 clutches, we started a new clone from the stock population. Mortality and clutch size measurements were used in a life-table analysis to estimate *r*.

### Trait measurements

We measured both morphological and life-history traits. Morphological traits were measured based on photos we took every other day of every individual. We used ImageJ software (Abràmoff et al. 2004) and a stage micrometer to measure three morphological traits at each time step based on photos. These traits were tail spine length (Havel and Dodson 1984, Ebert 1991), body size not including tail spine (Ebert et al. 1993), and head width (Havel and Dodson 1984). We also estimated three life history traits: clutch size (Spitze 1991) from counts of live born offspring, body size at first reproduction (Spitze 1991) from photos, and intrinsic population growth rate *r* (Dodson & Havel 1988) from a life-table analysis. To calculate *r*, we constructed a standard life table (Stearns 1992) for each population based on the reproduction and survivorship of individually raised clones. We estimated population-level level *l_x_* (stage specific survivorship) and *m_x_* (stage specific fecundity). We then used the *optim()* function in R 2.10 (Team 2012) to solve the Euler-Lotka equation (Stearns 1992) for *r*:

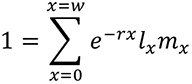

### Covariate measurement

We measured both biotic and abiotic covariates to test against *Daphnia* traits in a structural equation model (SEM). The abiotic variables measured weekly each summer were: pH, conductivity, dissolved O_2_, air temperature, water temperature, and light availability. We examined these variables to confirm that their distributions were stationary through time, and then calculated a single average for each pond across weekly samples over three years. We also collected soil cores from each pond to assess the potential effects of different allochthonous inputs of leaf litter. These cores were dried, sorted by tree species, and weighed. We used the cumulative weight for each tree species as a predictor variable. Finally, we measured the average predator density for all taxa that preyed on *D. pulex*. Predators were sampled with two cross sectional sweeps with a 10.2 × x 15.2 cm dip-net. We calculated the mean logarithm of predator abundance by summing the total number of predators on *D. pulex* from each sampling period and then taking the mean of the natural log of these abundances. These values were then averaged across each year to give mean log predator abundance for the duration of the study.

### Community composition

Twenty-seven different genera of aquatic animals were observed in the ponds over the course of three years. These taxa were almost all arthropods except for two anuran species: the green frog (*Rana clamitans*) and the wood frog (*Rana sylvatica*). The typical pond community consisted of four to ten different genera at any one sampling period. Three taxa of zooplankton other than *Daphnia pulex* were commonly observed*: Ceriodaphnia spp.,* Cyclopoid copepods, and Podocopid ostracods. Non-predatory taxa were either Diptera (larvae of mosquitoes, non-biting midges, or Dixid midges) or adult Coleoptera (family Hydrophilidae). The most common *Daphnia* predators were phantom midge larva (*Chaoborus spp*.), predaceous diving beetles (Dytiscidae: *Agabus spp* and *Accilius spp.*), and dragon fly nymphs (genus *Anax*), all of which are known to feed on *Daphnia pulex* (Kehl and Dettner 2003). We excluded water striders (genus *Gerris*), and Megaloptera larvae (Family Corydalidae, genus *Chauliodes*) from analysis as potential predators because we did not have evidence they specifically consume *Daphnia*.

### Statistical analysis

To test for direct effects of treatments on life history and morphology, we used a mixed model with nested random factors in R 2.14 (R Core Team 2012) with water loss rate and drought severity as fixed effects, and clone nested within isofemale line nested within pond as random effects (n = 108). Because the New England populations of *D. pulex* are obligately parthenogenic (Hebert and Finston 2001), we also used mixed models to partition phenotypic variance from the nested clonal design (Conner and Hartl 2004) to calculate broad-sense heritabilities. If experimental populations consisted of only one clone, there would be zero additive genetic variation and a heritability of zero. Population level traits such as *r* and traits such as clutch size and somatic growth rate for which sample sizes were not fully balanced due to mortality could not be analyzed with a mixed model. In these cases, we calculated pond level averages for each trait, treated each pond as replicate, and performed a two factor ANOVA (n = 12), with water loss rate and drought severity as the two crossed treatments. We also tested all traits against all measured covariates using linear regression, and tested for significant relationships between covariates and treatments using ANOVA. When we detected significant relationships among traits, treatments and covariates, we used a structural equation model (SEM) to tease apart indirect and direct relationships (Shipley 2004). Because of small sample sizes and the need for a continuous predictor variable for SEM analysis, we created a single continuous predictor variable from the sequential measurements of pond depth from each census. We calculated a pond coefficient of variation (C.V.) based on the weekly measurements of pond depth (C.V. = σ/µ, where σ and µ were estimated from eq. 6 and 7 in Ives et al. 2003 for a univariate time series).

In a full factorial ANOVA, water loss rate and drought severity accounted for 99% of the variation in pond C.V. (44% attributable to water loss rate, 55% to drought severity, and the remainder to error and interactions). Thus, pond C.V. effectively captured the variation imposed by the two experimental treatments as a single continuous variable for fitting an SEM. The SEM analysis consisted of 18 different models, all specified and run in the software package lavaan (Rosseel 2012). We tested a null model that included only traits, and a model with just pond C.V. directly affecting the traits. The remaining 16 models included one of the 8 possible covariates in two variations. The first model variation included pond C.V. directly linked only to a covariate; therefore it could only indirectly affect the response variable. The second model included a direct link to both the covariate and the two response variables. Multiple model selection criteria (BIC, AIC and AICc) were used to determine the best-fitting model (Burnham and Anderson 2010).

## Results

### Direct response of traits to experimental treatments

Of the seven measured traits four showed at least one significant response based on ANOVA and mixed models: tail spine length at first instar (hereafter tail spine length), population growth rate (*r),* body size at first reproduction, and average clutch size (Table 1). Tail spine length and *r* both responded significantly to water loss rate and drought severity treatments (Figure 1). Tail spine length and *r* were largest in treatments with low water loss rates and low drought intensity (simulation of current climate conditions). We quantified reductions in tail spine length and *r* as 1 − *x̄*_*Future*_/*x̄*_*Current*_, one minus the trait mean measured under future climate scenarios (high water loss rate, high drought severity) divided by the trait mean measured under current climate scenarios (low water loss rate, low drought severity). Tail spine length was 15% shorter in high water loss rate and high drought treatments (simulation of future climate conditions; Figure 1, Figure 2). Similarly, *r* was 18% lower in the high water loss rate and high drought treatments (Figure 1). We calculated broad-sense heritablitilies for tail spine to ensure that ponds had existing additive genetic variance. Broad-sense heritabilities for tail spine length ranged from 18.5% to 74.8% with a mean of 44.3%, which is comparable to other published estimates for the heritability of morphological characters in *Daphnia* (Ebert et al. 1993). Size at first reproduction was significantly lower in high drought severity treatments (*x̄* = 1.26 *mm*, n = 12, d.f. = 1, F= 6, p < 0.05) compared to low drought severity treatments (*x̄* = 1.38 *mm*). The average clutch size was significantly larger in high water level treatments (*x̄* = 11.3 daughters / clutch, n, = 12, d.f. = 1, F = 9.75, p < 0.05) compared to low water level treatments (*x̄* = 9.5 daughters / clutch). All other traits showed no significant response.

**Figure 1.**
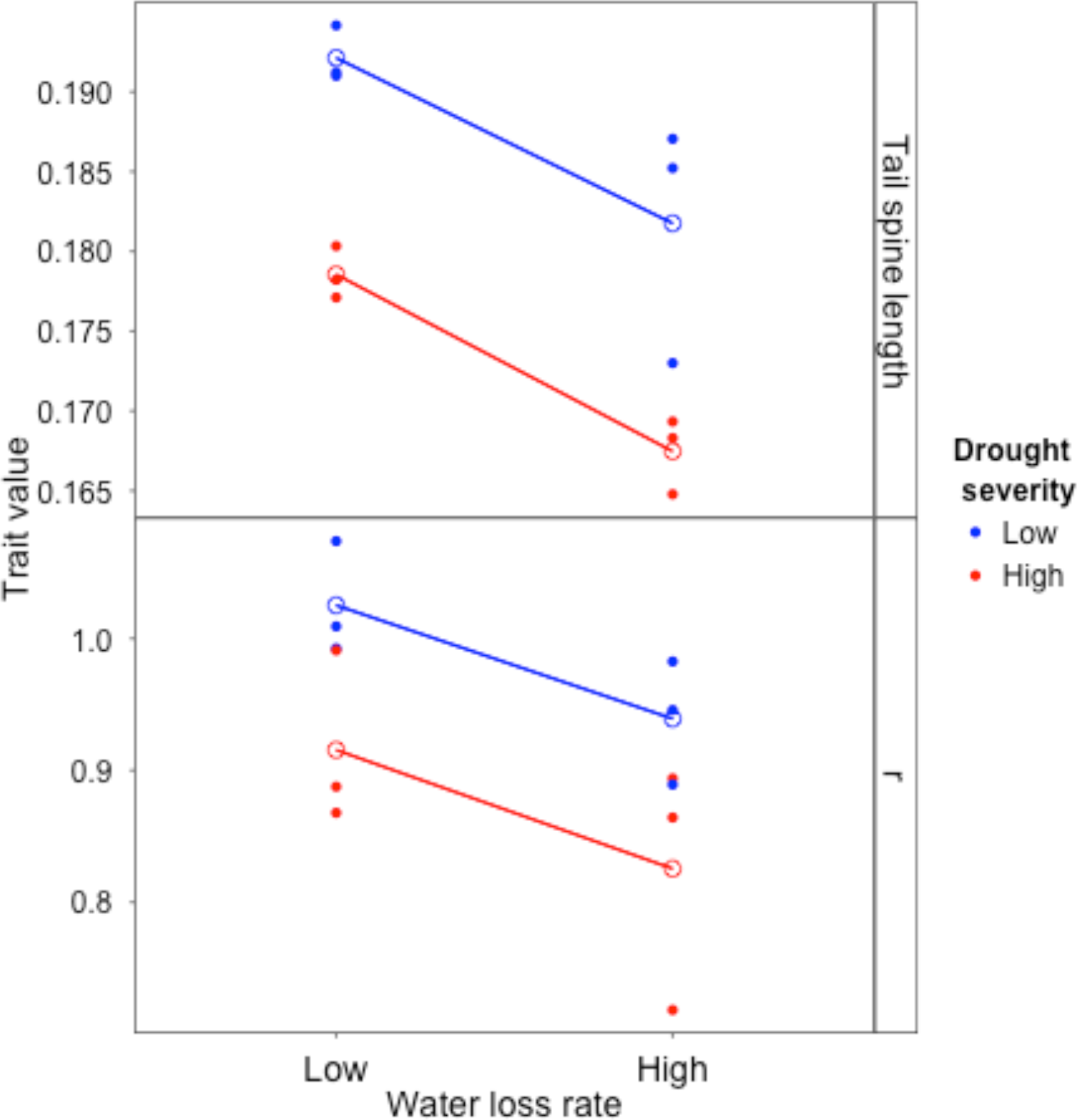
Evolutionary changes in morphology (tail spine length) and life history (*r*, the intrinsic rate of increase) of *Daphnia pulex* in response to experimental manipulations of pond hydrology representing different scenarios of climate change. Each solid point is the response of the population from a single pond in an orthogonal 2-factor field experiment in which water loss rate and drought severity were manipulated for 3 consecutive years. Open circles represent the grand mean of each treatment combination. The longest tail spines and highest *r* values were measured in ponds that mimic current climate (low water loss rate and low drought severity) and have the greatest number of predators.

**Figure 2.**
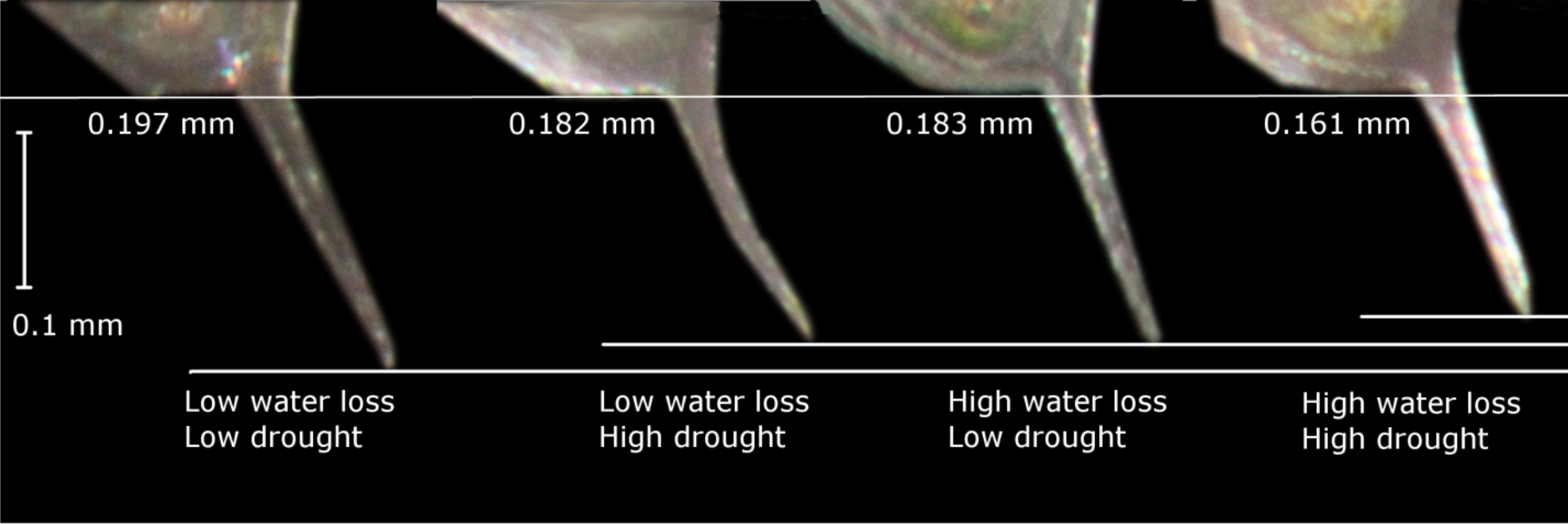
Representative tail spine lengths of first instar *D. pulex* collected from four experimental climate change treatments, and reared for 3 clonal generations in common garden conditions. Tail spine length exhibited an additive response to the climate change treatments and is greatest in the low water-loss-rate, low drought-severity treatment combination.

**Table 1.** F-ratios of ANOVA models for seven morphological and life history traits.

| *Factor* | *Spine*<br>*Length* | *Body*<br>*size* | *Head*<br>*width* | *Clutch*<br>*size* | *Growth*<br>*rate* | *Size at first*<br>*reproduction* | *r* |
| --- | --- | --- | --- | --- | --- | --- | --- |
| Water loss rate | **36.3** | 0.51 | 0.04 | **9.75** | 0.26 | 0.94 | **5.38** |
| Drought severity | **42.1** | 0.003 | 0.13 | 2.53 | 0.18 | **5.99** | **8.68** |
| WLR * DS | 0.42 | 0.47 | 0.42 | **7.52** | 0.07 | 0.00 | 0.00 |

Table 1. F-ratios from a two-way ANOVA with interactions for pond level averages of all measured traits; significant effects (*p* < 0.05) are in bold, with 1 degree of freedom for each model term and 8 residual degrees of freedom.

### Response of traits to measured covariates

Both tail spine length (R^2^ = 77%, p < 0.05) and *r* (R^2^ = 69%, p < 0.05) were significantly correlated with mean predator abundance (Figure 3A). No other *Daphnia* traits were significantly correlated with predator abundance. No traits were significantly correlated with any of the other environmental covariates (Figure 4). The most parsimonious SEM model was one that included predators having a direct effect on traits, mediated through pond C.V. The second most parsimonious model included direct effects of pond C.V. on traits, but those direct effects were not significant (Figure 3B, ΔBIC > 2). The next closest models in BIC value included pH or did not include any covariate. These models all had substantially larger ΔBIC values (ΔBIC > 22), implying strong support for the best-fitting model that included direct effects of predators and an indirect effect of pond C.V. (Table 1).

**Figure 3.**
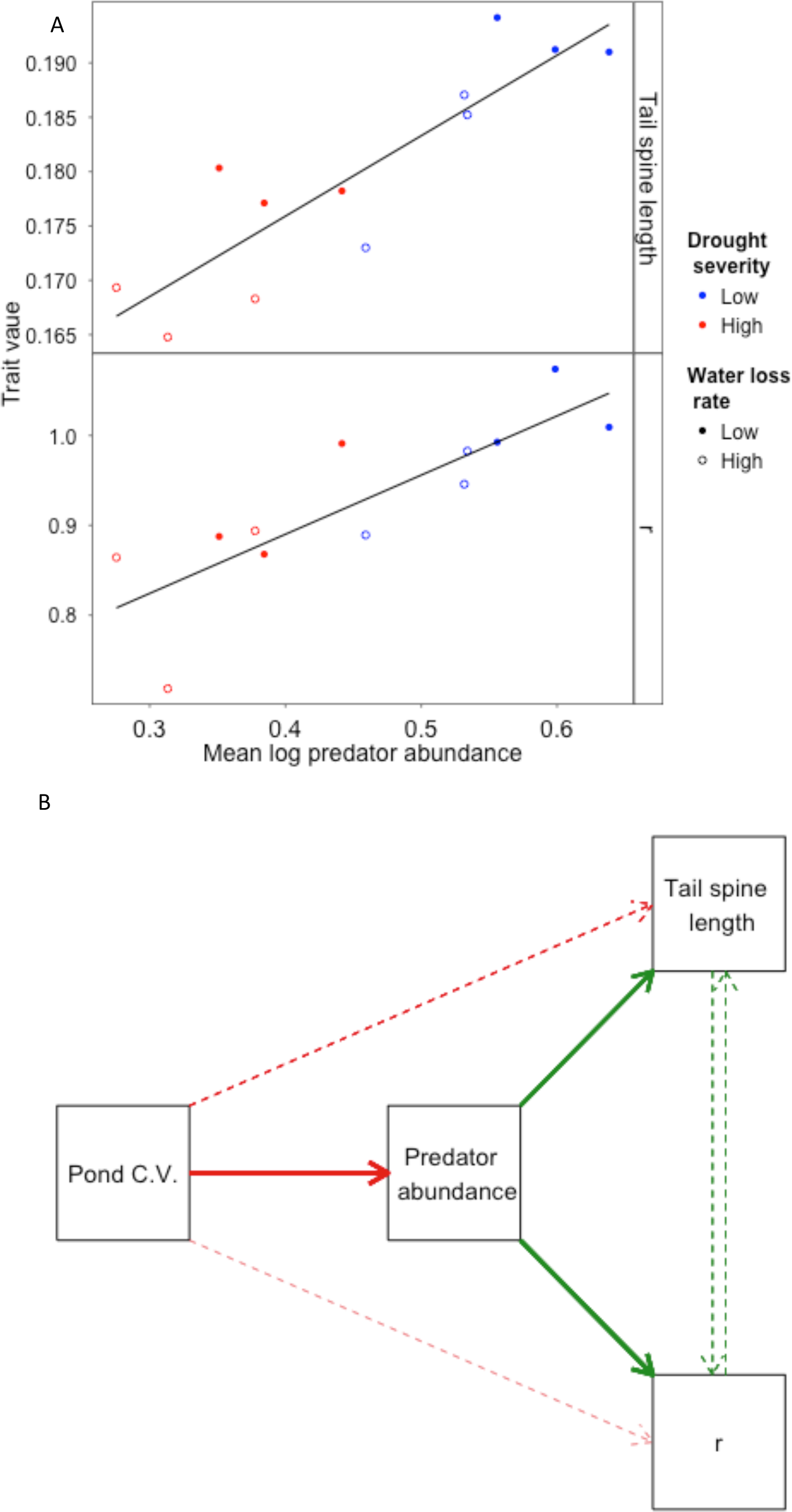
A) Linear regression of average *D. pulex* tail spine length (mm) and intrinsic rate of increase (*r*; individuals/individual•day) versus daily average log predator abundance (R^2^ = 0.77, *P* = 0.0001 for tail spine and R^2^ = 0.69, *P* = 0.0008 for *r*). B.) The best-supported SEM analysis model for evolutionary changes in tail spine length and intrinsic rate of increase (*r*) of *Daphnia pulex* populations exposed to a 3-year climate change experiment. Solid arrows represent statistically significant SEMs; dashed arrows represent non-significant SEMs. The width of each arrow is proportional to the standardized model coefficient (red arrows = negative effects, green arrows = positive effects). Pond C.V. is a continuous composite continuous variable based on the experimental treatments.

**Figure 4.**
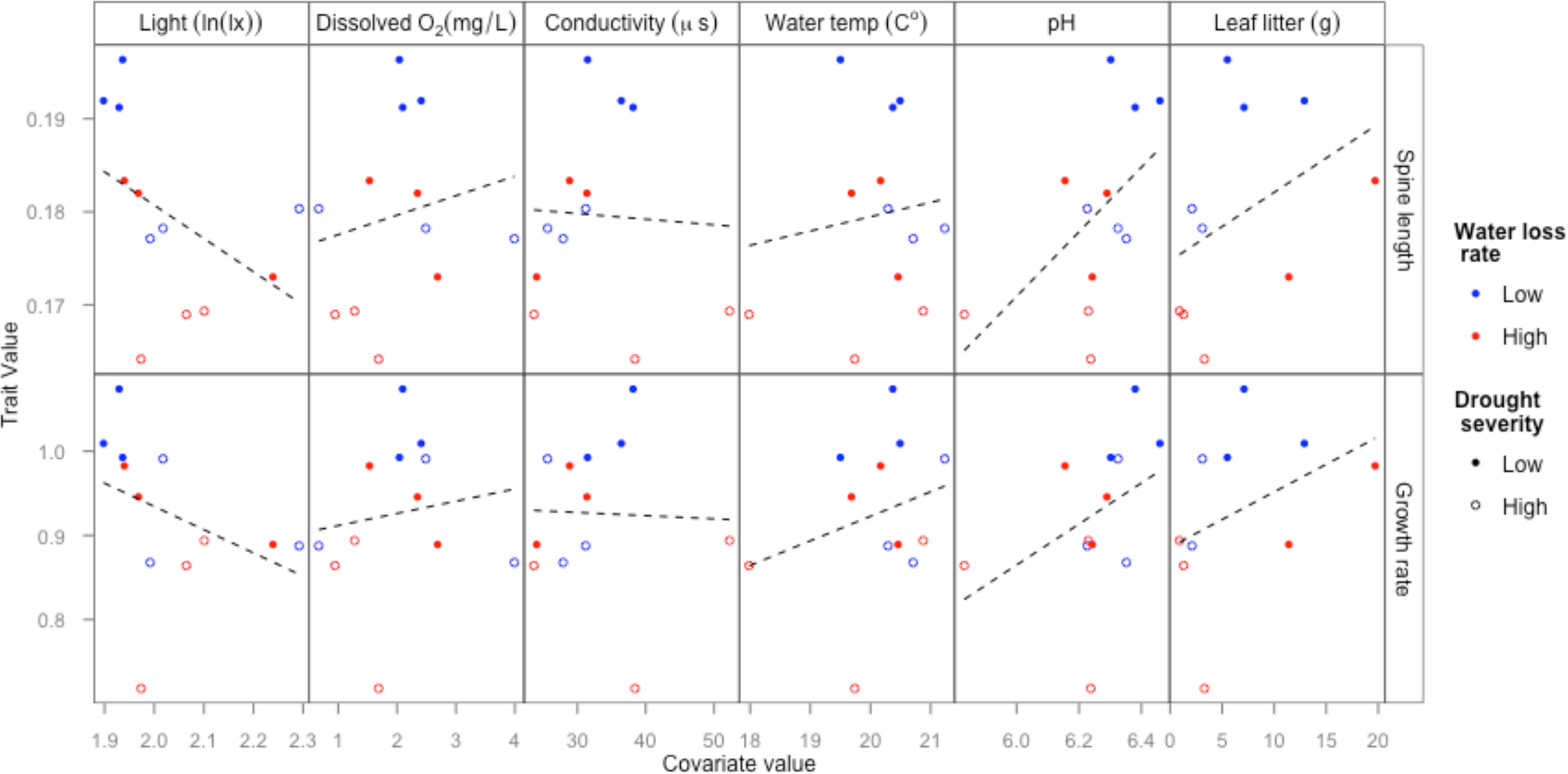
Spine length and r plotted against other potential abiotic and biotic covariates. Each point represents the average for a single experimental pond (n = 12)/ Covariate measures are pond level averages calculated over the 3 years of treatment application. No significant relationship was found among any other variables, indicated by dashed lines. Open and closed circles indicate water loss rate, blue and red colors indicate drought severity.

## Discussion

Although the importance of eco-evolutionary feedbacks is becoming increasingly recognized (Ellner et al. 2011, Urban et al. 2012a, Walsh et al. 2012), ecologists have often neglected rapid evolutionary responses and have mostly emphasized simple ecological responses of assemblages to abiotic conditions that are expected with future climate change. Here we have provided evidence from a multi-generation field experiment that abiotic climate change can restructure ecological communities via shifts in the abundance of predators, and that the new community structure is itself an evolutionary selective force. *Daphnia* populations in experimental ponds responded strongly to predator abundance (Figure 1, Figure 2), but not to measured changes in abiotic factors associated with simulated climate change (Figure 4))

Although some traits showed a clear selective response to predation (Figure 1), there was no significant response in somatic growth rate or body size (Table 1), which some other investigators have observed (Spitze 1991). The *Daphnia* in this experiment were all originally collected from the same vernal pond, and perhaps there was little additive genetic variance due to genetic isolation of this single source (Haag et al. 2006). However strong founder effects caused by the presence of only a few clones colonizing a pond (Allen et al. 2010) seem unlikely in this experiment because the ponds were seeded with ∼150 individuals each, and because the system was always open to external colonization during the 3 years of the experiment. *Daphnia* can respond rapidly to selection (Hairston et al. 1999), which is consistent with the strong effects measured in response to experimental alterations of hydroperiod (Figures 1 and 2), and traits responded in the direction consistent with previous experimental studies (Spitze 1991) and with the predictions of life history theory (Taylor and Gabriel 1992).

*D. pulex* with longer tail spines are less vulnerable to predation (Havel and Dodson 1984, Dodson and Havel 1988), possibly due to increases in predator handling time or changes in prey buoyancy (Lüning 1992). Predator abundances are predicted to decrease as habitat variability increases because of habitat preferences in colonization and longer development times (Schneider 1997). In response to climate change manipulations, the evolved trait changes measured in *D. pulex* were comparable to those found in laboratory experiments on the response of *D. pulex* to predators (Spitze 1991, Lüning 1992). In those earlier studies, first-instar tail spines of *D. pulex* populations that were exposed to multiple predators in the laboratory increased in length by 12% (Spitze 1991). In some studies, increases in tail spine length are an induced, phenotypically plastic, response to predators (Lüning 1992). However, in this study, *Daphnia* collected from the field were reared through multiple generations in laboratory conditions in the complete absence of predators or water-borne chemical cues associated with predators. Moreover, neck teeth in *Daphnia pulex* are the characteristic sign of a phenotypically plastic response to kairomones (Riessen 1999), but they were never observed in our laboratory-reared populations.

Life-history theory predicts that, in the presence of predators that feed selectively on small size classes (Spitze et al. 1991), prey populations should evolve delayed reproduction, increased investment in early somatic growth, and greater fecundity at later instars. These life-history shifts may lead to increases in *r* (Brett 1992, Taylor and Gabriel 1992). Population growth rates were greatest in ponds with the highest predator densities. Results of the common garden experiment were also consistent with other predictions of life-history theory and previous *Daphnia* laboratory studies: with decreasing predation pressure, body size at first clutch decreased, and clutch sizes of older age classes decreased (Spitze 1991, Brett 1992).

Two lines of evidence suggest that the morphological and life history changes in *Daphnia* lineages from different ponds reflect evolutionary responses to predators, rather than evolutionary responses to altered abiotic conditions. First, none of the measured abiotic variables in each pond (pH, conductivity, dissolved O_2_, water temperature, and light availability) was correlated with *Daphnia* spine length and *r* (Figure 4). In contrast, average predator abundance (the 3-year average of weekly measurements of the logarithm of the abundance of all predatory taxa in a pond) was highly correlated and explained most of the variation among ponds in tail spine length (77%) and *r* (69%; Figure 3A). Second, the best-fitting SEM model included direct effects of predators but did not include direct effects of the experimental treatments on the response variables (Figure 3B). This SEM model fit the data substantially better than an alternative model that included only treatment effects and no predator covariate (ΔBIC = 33) and better than a null model that included only correlations between the response variables tail spine length and *r* (ΔBIC = 80). Collectively, these analyses suggest that observed trait differences are genetically based evolutionary responses reflecting altered interactions with predators, rather than direct responses to altered abiotic conditions.

Collectively, our results suggest that climate change can trigger a cascading response in which both altered abiotic conditions and species interactions can affect populations through ecological and evolutionary pathways. The indirect effects of altered species interactions such as predation, parasitism, and competition may be just as important as the direct effects of altered temperature, precipitation, and concentrations of greenhouse gasses on the response of species and populations to global climate change.

## Acknowledgements

We thank A. Brody, S. Helms-Cahan, G. Gilchrist, E. Wolkovich, and S. Lavergne for comments on the manuscript; D. Brynn and D. Tobi for logistic support at the UVM Jericho Research Forest; A. Brody for laboratory space, and C. Love for laboratory assistance. This project was supported by a grant from VT EPSCoR NSF (EPS #0701410) and an NSF DDIG (DEB-0909359) awarded to the authors.

